# Argonaute-siRNA loading via the RNA-binding protein RDE-4 in *C. elegans*

**DOI:** 10.1101/2025.05.06.652520

**Authors:** Thiago L. Knittel, Brooke E. Montgomery, Reese A. Sprister, Colin N. Magelky, Margaret J. Smith, Maritza Soto-Ojeda, Melissa Guthrie, Carolyn M. Phillips, Taiowa A. Montgomery

## Abstract

Small RNAs, such as microRNAs (miRNAs) and small interfering RNAs (siRNAs), associate with Argonaute proteins to control gene expression, impacting a wide range of cellular processes, such as antiviral defense, transposon silencing, and development^1^. Plants and animals typically have several classes of small RNAs, along with multiple Argonautes^2^. These Argonautes often confer distinct functionality to the various classes of small RNAs^3^. But how small RNAs are selectively loaded into the appropriate Argonaute is not well understood. miRNAs and siRNAs are typically generated from double-stranded RNA (dsRNA) precursors by the endoribonuclease Dicer^4^. siRNAs are often processed from fully base-paired precursors derived from various endogenous and exogenous sources, whereas miRNAs typically originate from genetically encoded partially base-paired hairpins^1^. In *C. elegans*, Dicer/DCR-1 processing of siRNAs and a related small RNA class, known as 26G- RNAs, is mediated by the dsRNA-binding protein RDE-4^5-7^. Here, we show that RDE-4 also facilitates loading of siRNAs (but not miRNAs) into the Argonaute RDE-1, but not into ALG-1, and loading of 26G-RNAs into the Argonaute ERGO-1. Although we do not find evidence that ALG-3/4 associated 26G-RNAs require RDE-4 for Argonaute loading, their levels are strongly reduced in *rde-4* mutants indicating that RDE-4 is broadly required for their formation or stability. Our findings reveal a role for RDE-4 as a critical determinant of small RNA loading specificity and provide insight into the mechanisms by which small RNAs are selectively paired with their corresponding Argonautes.

## RESULTS AND DISCUSSION

### Requirement of RDE-4 in endogenous and exogenous small RNA pathways

RDE-4 was first identified for its role in exogenous RNA interference (RNAi) over two decades ago^8^. It interacts with dsRNA and resides in a complex containing DCR-1, and the RIG-I ortholog DRH-1, which processes dsRNA into siRNAs^8-17^. RDE-4 has been implicated in antiviral defense, exogenous RNAi, and multiple endogenous small RNA pathways^18-20^. The protein functions analogously to Loqs-PD in *Drosophila*, which is also required for processing dsRNA into siRNAs^21-26^. In *Drosophila,* another dsRNA-binding protein called R2D2 is necessary for assembling siRNAs into the Argonaute Ago2 silencing complex, however, an equivalent protein has not been identified in *C. elegans*^27-30^. Loqs-PD, R2D2, and RDE-4 each contain two dsRNA binding domains that adopt α-β-β-β-α folds (Figure 1A). Based on this structural similarity, we hypothesized that RDE-4 performs the functions of both Loqs-PD and R2D2. The potential role of RDE-4 in connecting dsRNA processing and Argonaute loading was initially proposed by Liu et al.^27^ based on its sequence similarity to R2D2. However, to our knowledge, this has not been experimentally tested. We reasoned that if RDE-4 facilitates Argonaute-siRNA interactions, siRNAs would be depleted in Argonaute co-immunoprecipitates (co-IPs) from *rde-4* mutants. However, because a comprehensive analysis of small RNAs in *rde-4* mutants was lacking, it was unclear if sufficient levels of residual siRNAs are present in *rde-4* mutants to assess their association with Argonautes. Thus, we first assessed the global impact of loss of *rde-4* on small RNAs using small RNA high-throughput sequencing (sRNA-seq).

**Figure 1.**
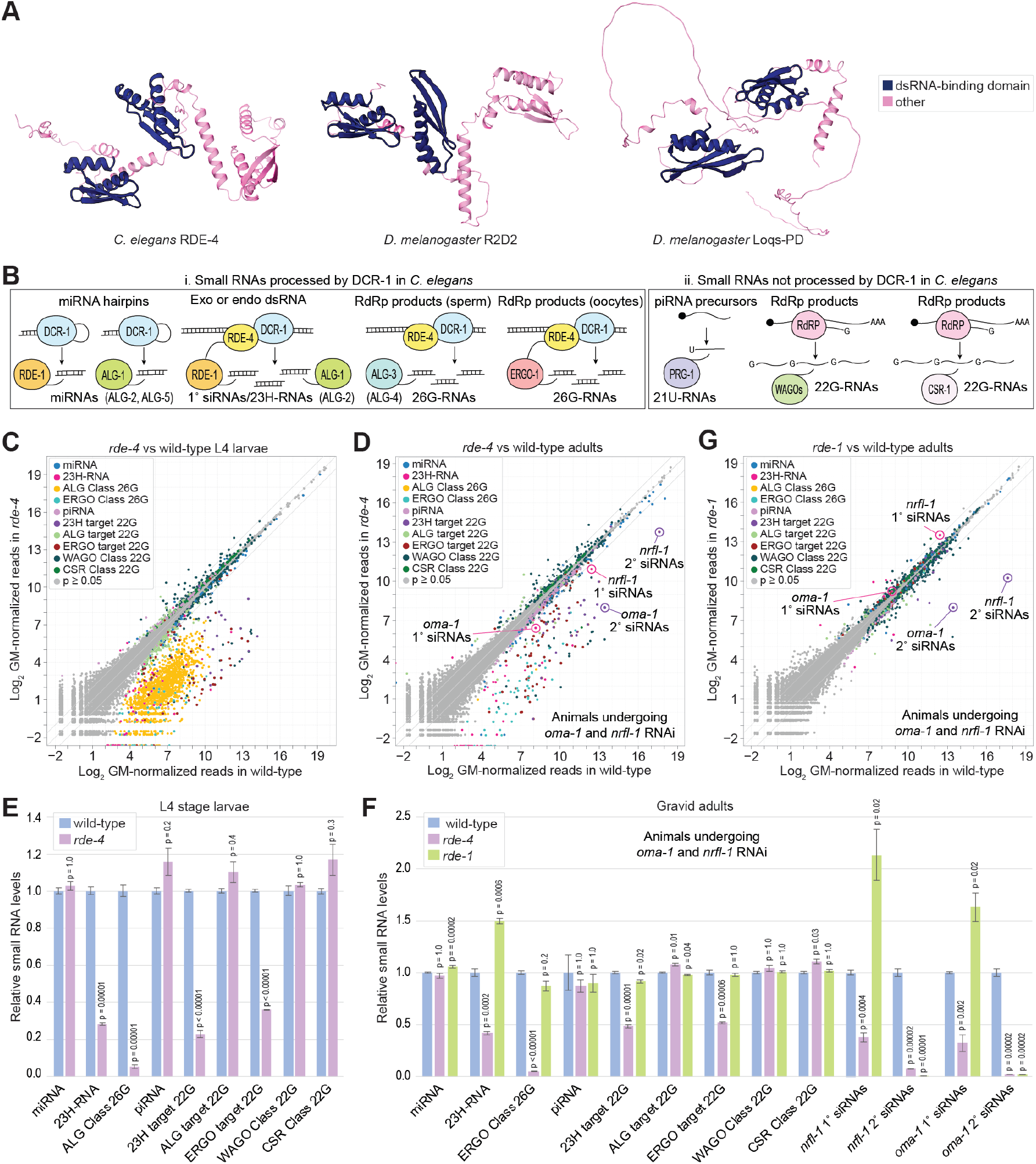
Requirement of *rde-4* across different small RNA pathways. **(A)** AlphaFold3-predicted structures of *C. elegans* RDE-4, and *Drosophila* R2D2 and Loqs-PD. dsRNA binding domains are highlighted. **(B)** Schematic overview of *C. elegans* small RNA classes, indicating their Argonaute-binding partners, DCR-1 processing dependency, and whether they are synthesized by RdRPs. Exo, exogenous. Endo, endogenous. **(C, D)** Scatter plots showing individual small RNA features as the average log_2_ GM-normalized sRNA-seq reads in wild-type (x-axis) and *rde-4* mutant (y-axis) animals. Small RNA classes are color-coded. Exogenous siRNAs mapping to *nrfl-1* and *oma-1* are circled. Samples are from L4 stage larvae (C) or gravid adults (D). RNA was treated with RppH. n=3 biological replicates. **(E)** Relative rpm-normalized abundance of small RNA classes in wild-type and *rde-4* mutant L4 larvae (data as in C). Error bars represent standard deviation (SD) from the mean. *p*-values were calculated using two-sample t-tests. **(F)** Same as in (E), but with gravid adult wild-type, and *rde-4* and *rde-1* mutant animals (data as in D). Error bars represent SD from the mean. *p*-values were calculated using two-sample t-tests. **(G)** Scatter plot as in (C-D), comparing wild-type (x-axis) and *rde-1* mutant (y-axis) gravid adults. Small RNA classes are color-coded; *nrfl-1* and *oma-1* siRNAs are circled. n=3 biological replicates (data also in F).

*C. elegans* contains several classes of small RNA, some of which are directly processed by DCR-1 and others that are not (Figure 1B)^31^. miRNAs were not affected in *rde-4* mutants in either L4 larval stage or gravid adult animals (Figures 1C-1F; Tables S1-S2). In contrast endogenous canonical siRNAs, also called 23H-RNAs, were modestly depleted in *rde-4* mutants (Figures 1C-1F; Tables S1-S2)^32^. The adult stage animals used in these sRNA-seq experiments were subjected to exogenous RNAi targeting both *nrfl-1* and *oma-1*, genes that can be depleted without leading to obvious developmental defects^33^. Exogenous siRNAs derived from the *nrfl-1* and *oma-1* dsRNA administered to the animals were also modestly reduced in *rde-4* mutants (Figures 1D and 1F; Tables S1-S2).

*C. elegans* contains two classes of non-canonical siRNAs that are 26-nucleotides (nt) long and possess a 5’G. One class is produced in sperm development during the L4 stage and bind the Argonautes ALG-3 and ALG-4 and the other is produced from a largely non-overlapping set of genes during oocyte formation, mostly in the adult stage, and bind the Argonaute ERGO-1 (Figure 1B). These so-called 26G-RNAs are produced from RNA-dependent RNA polymerase (RdRP) products, which are processed by DCR-1, but mechanistic details are lacking^5,6,34-36^. Both ALG-3/4 and ERGO-1 classes of 26G-RNAs, were strongly depleted in *rde-4* mutants (Figures 1C-1F; Tables S1-S2). Exogenous siRNAs, 23H-RNAs, and ERGO-1 class 26G-RNAs trigger the production of secondary small RNAs called 22G-RNAs, which are not made by DCR-1 (Figure 1B)^5,6,37,38^. These secondary 22G-RNAs were also depleted in *rde-4* mutants (Figures 1C-1F; Tables S1-S2). However, most 22G-RNAs are not dependent on 1° siRNAs for their formation and these small RNAs, as well as DCR-1-independent piwi-interacting RNAs (piRNAs), were not depleted in *rde-4* mutants (Figures 1C-1F; Tables S1-S2)^31^. Thus, RDE-4 is specific to DCR-1-dependent siRNA pathways, although the extent to which it is required for small RNA formation or stability differs between these pathways.

Both exogenous siRNAs and 23H-RNAs associate with the Argonautes RDE-1, ALG-1, and ALG-2, but are particularly enriched in RDE-1^32^. Despite the strong bias of RDE-1 for canonical siRNAs their levels were not reduced in *rde-1* mutants, indicating that this association is not important for siRNA stability (Figures 1F-1G; Table S2)^32^. However, the secondary 22G-RNAs generated downstream are depleted, as their production relies on RDE-1, which is not compensated for by ALG-1 or ALG-2 (Figures 1F-1G; Table S2)^8,32^. The substantial presence of DCR-1-dependent siRNAs remaining in *rde-4* mutants enabled us to test our hypothesis that RDE-4 plays a role in Argonaute loading.

### RDE-4 facilitates loading of exogenous siRNAs into RDE-1

To assess whether RDE-4 promotes the association of exogenous siRNAs with RDE-1, we co-IP’d GFP::RDE-1 from either *rde-4+/+* or *rde-4-/-* animals undergoing RNAi against *nrfl-1.* We sequenced the associated small RNAs and compared them to those present in the input cell lysates. In both cell lysates and co-IP fractions from *rde-4+/+* and *rde-4-/-* animals, *nrfl-1* 1° siRNAs displayed similar size and 5’ nucleotide distribution profiles, with an enrichment of 23-nt reads (Figures 2A-2B). Consistent with our findings above, we observed a 1.7-fold decrease in *nrfl-1* 1° siRNA levels in cell lysates from *rde-4-/-* animals, confirming the requirement of RDE-4 for siRNA formation or stability (Figure 2A). However, in GFP::RDE-1 co-IP fractions, we observed a 42-fold reduction in *nrfl-1* 1° siRNA reads in *rde-4-/-* animals, which is nearly 25-fold greater than the reduction observed in cell lysates (Figure 2B). Additionally, *nrfl-1* 1° siRNA reads in GFP::RDE-1 co-IPs from *rde-4+/+* animals were enriched 56.5-fold relative to the input cell lysates, whereas enrichment was reduced to only 1.3-fold in *rde-4-/-* animals (Figure 2C). This indicates that a reduction in siRNA formation or stability was not solely responsible for the lower levels of siRNAs in GFP::RDE-1 co-IPs from *rde-4* mutant animals.

**Figure 2.**
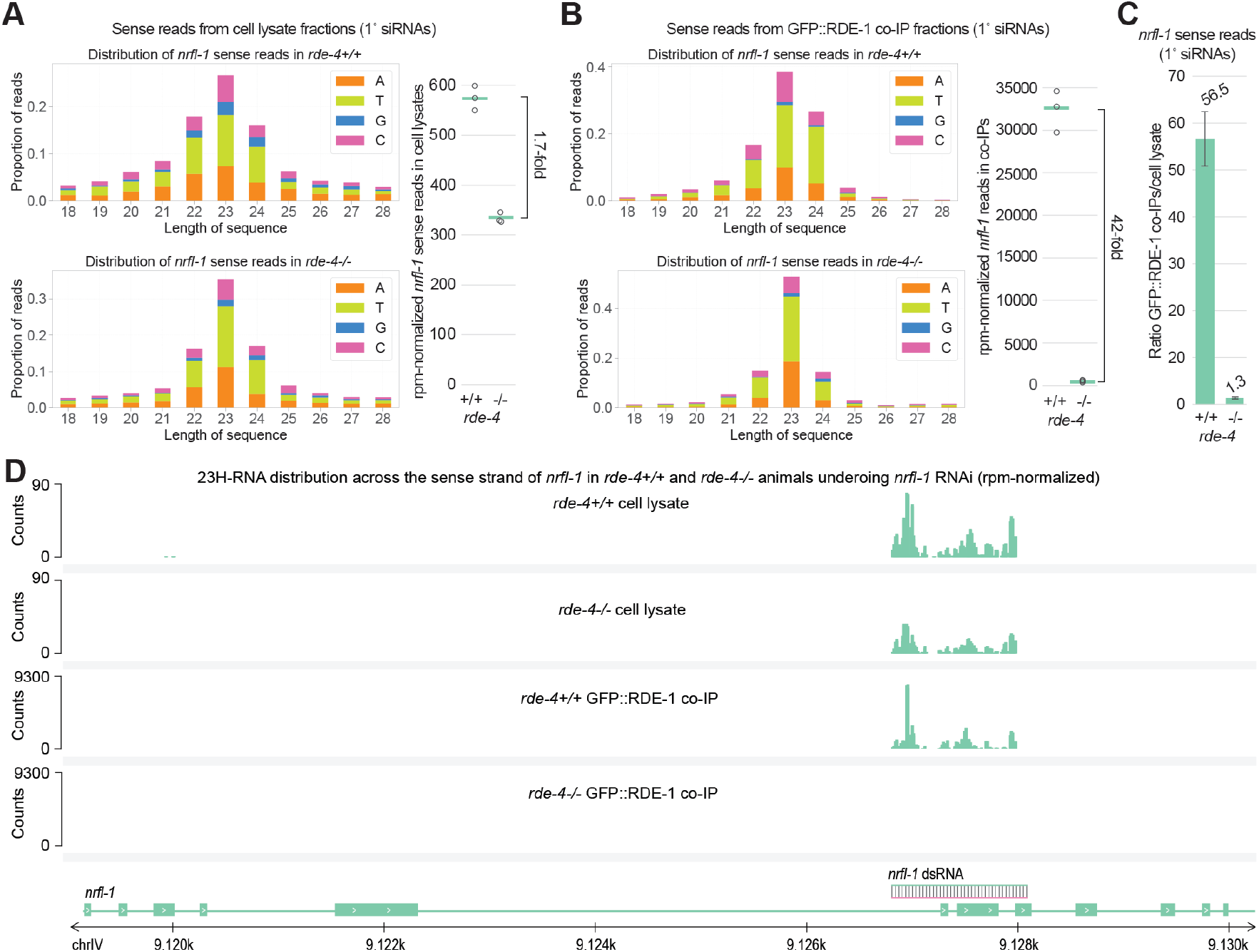
RDE-4 facilitates association of exogenous siRNAs with RDE-1. **(A-B)** Size distribution and 5′-nucleotide identity of *nrfl-1* sense siRNAs from cell lysates (**A**) and GFP::RDE-1 co-IPs (**B**) from *rde-4+/+* or *rde-4-/-* mutant gravid adults. One representative of 3 biological replicates is shown. The plots on the right show the average rpm-normalized counts for *nrfl-1* sense siRNA reads from three biological replicates. **(C)** Enrichment of total *nrfl-1* sense exogenous siRNA reads in GFP::RDE-1 co-IPs relative to corresponding cell lysates from *rde-4+/+* or *rde-4-/-* mutant gravid adults (data as in A-B). Error bars represent SD from the mean of three biological replicates. **(D)** rpm-normalized small RNA-seq read distribution across the sense strand of *nrfl-1* from cell lysates and GFP::RDE-1 co-IPs from *rde-4+/+* or *rde-4-/-* animals.

siRNAs were modestly and uniformly depleted across *nrfl-1* dsRNA in cell lysates from *rde-4-/-* animals (Figure 2D). The strong enrichment observed in GFP::RDE-1 co-IPs from *rde-4+/+* animals was also uniformly lost in *rde-4-/-* mutants, demonstrating that RDE-4 is broadly required for optimal siRNA association with RDE-1 (Figure 2D). Similar results were obtained in an independent experiment utilizing animals treated with both *nrlf-1* and *oma-1* RNAi (Figures S1A-S1F). That siRNAs still associate with RDE-1 in the absence of *rde-4*, albeit at much lower levels, likely explains why the RNAi defects of *rde-4* mutants can be overcome by injecting high levels of dsRNA^39^.

### Endogenous siRNAs require RDE-4 for optimal loading into RDE-1

We next tested whether RDE-4 facilitates loading of endogenous siRNAs (i.e. 23H-RNAs) into RDE-1. Individual 23H-RNAs were highly enriched in RDE-1 co-IPs relative to cell lysates, comparable to *nrfl-1* exogenous siRNAs, following geometric mean (GM)-based normalization (Figure 3A; Table S3). Numerous miRNAs were also enriched in GFP::RDE-1 co-IPs, but generally to a much lesser extent than 23H-RNAs (Figure 3A; Table S3). Strikingly, the enrichment of *nrfl-1* 1° siRNAs and 23H-RNAs was reduced to levels similar to miRNAs in *rde-4-/-* animals (Figure 3B; Table S3). The ratio of total 23H-RNA reads, normalized by million mapped reads (rpm), in GFP::RDE-1 co-IPs relative to cell lysates was reduced by ∼36-fold in *rde-4-/-* animals, while miRNA enrichment was unchanged (Figure 3C). Unlike GM normalization, rpm normalization does not account for variation in individual small RNAs across samples. While GM normalization more accurately identifies differences in individual small RNAs, it is generally less reliable when assessing total reads for a specific class of small RNAs, based on our experience.

**Figure 3.**
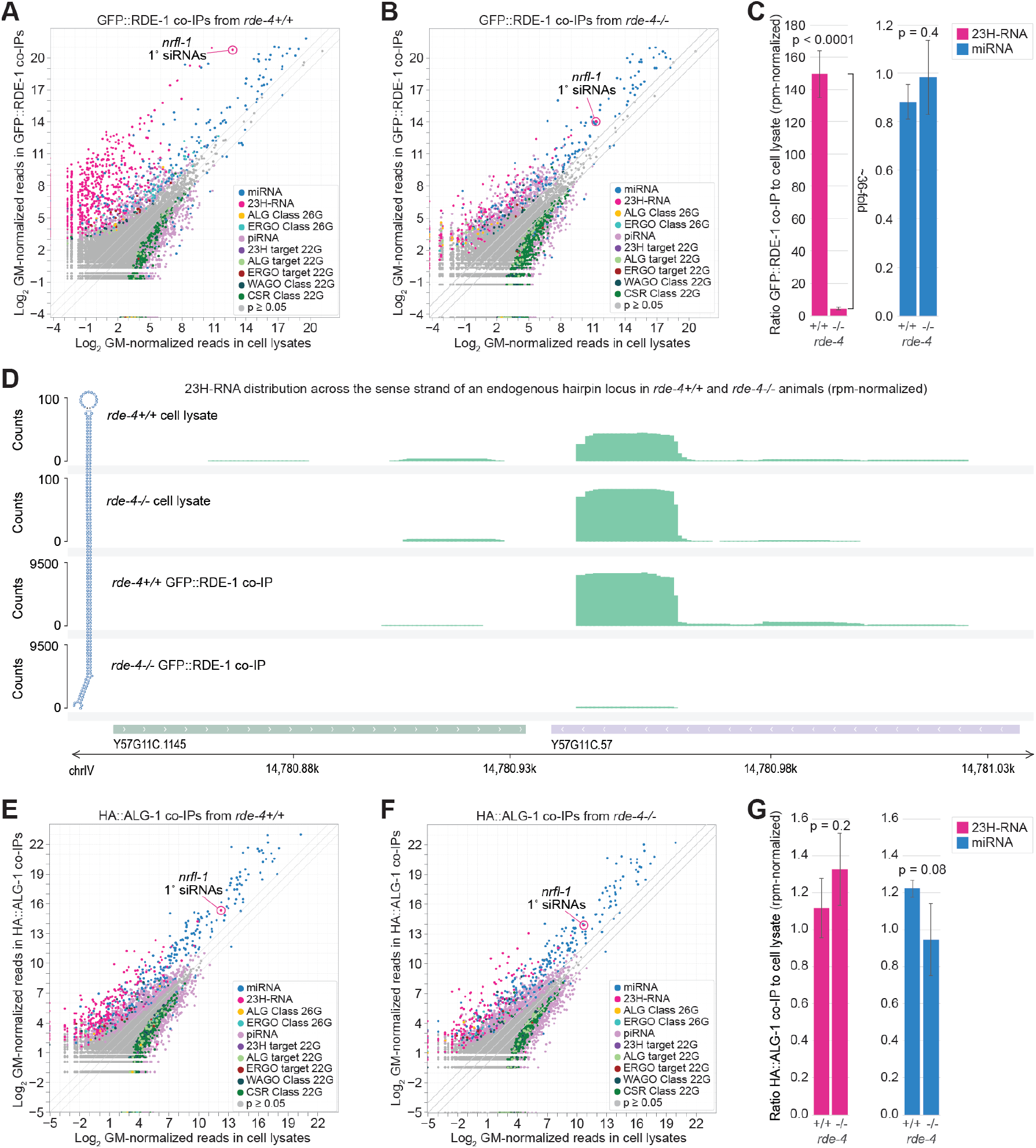
RDE-4 directs endogenous siRNA association with RDE-1 but not ALG-1. **(A-B)** Scatter plots showing individual small RNA features as the average log_2_ GM-normalized sRNA-seq reads in cell lysates (x-axis) and GFP::RDE-1 co-IPs (y-axis) from *rde-4+/+* (A) or *rde-4-/-* (B) animals undergoing *nrfl-1* RNAi. Small RNA classes are color-coded. n=3 biological replicates. **(C)** Average enrichment of total rpm-normalized 23H-RNA and miRNA reads in GFP::RDE-1 co-IP libraries relative to corresponding cell lysates (data as in A-B). *p*-values were calculated using two-sample t-tests. **(D)** rpm-normalized small RNA-seq read distribution across the sense strand of Y57G11C.57 from cell lysates and GFP::RDE-1 co-IPs from *rde-4+/+* or *rde-4-/-* animals. **(E-F)** Scatter plots as in (A-B) but comparing cell lysates (x-axis) and HA::ALG-1 co-IPs (y-axis). n=3 biological replicates. **(G)** Average enrichment of total rpm-normalized 23H-RNA and miRNA reads in HA::ALG-1 co-IPs relative to corresponding cell lysates (data as in E-F). *p*-values were calculated using two-sample t-tests.

The reduction in siRNA enrichment in GFP::RDE-1 co-IPs from *rde-4-/-* animals could be indirect, resulting from reduced siRNA abundances in cell lysates, such that the residual siRNAs are competed away from RDE-1. To test this possibility, we co-IP’d GFP::RDE-1 from animals undergoing RNAi against *dcr-1*, which, like loss of *rde-4*, resulted in a reduction in the levels of a representative endogenous 23H-RNA derived from the F43E2.6 locus, as determined by quantitative real-time PCR (qRT-PCR) (Figures S1G-S1H). Although the levels of the F43E2.6 23H-RNA were reduced, its enrichment in RDE-1 co-IPs was unchanged in *dcr-1* RNAi treated animals (Figure S1H). This suggests that impaired loading of siRNAs into RDE-1 in *rde-4* mutants is not due to reduced siRNA abundance. Furthermore, even 23H-RNAs that normally associate strongly with RDE-1 but that were not depleted in the cell lysates of *rde-4-/-* animals were nevertheless depleted in GFP::RDE-1 co-IPs from *rde-4-/-* animals (Figure 3D; Table S3).

We observed a similar loss of hyperenrichment of 23H-RNAs and exogenous *nrfl-1* and *oma-1* 1°siRNAs in GFP::RDE-1 co-IPs from *rde-4* mutants in an independent experiment (Figures S2A-S2B). We conclude that although not absolutely required for loading siRNAs into RDE-1, RDE-4 facilitates their preferential association over miRNAs. Because siRNAs are typically much less abundant than miRNAs, RDE-4 may be necessary to ensure that sufficient siRNA-RDE-1 complexes are formed during RNAi.

### RDE-4 does not promote ALG-1-siRNA interactions

Exogenous siRNAs and 23H-RNAs also associate with the major miRNA Argonautes ALG-1 and ALG-2^32,40,41^. We questioned whether RDE-4 is required for loading siRNAs into these Argonautes as well. To address this, we subjected small RNAs from HA::ALG-1 co-IPs and cell lysates from *rde-4+/+* and *rde-4-/-* animals treated with *nrfl-1* RNAi to sRNA-seq. miRNAs, *nrfl-1* exogenous siRNAs, and 23H-RNAs were all similarly enriched in HA::ALG-1 co-IPs compared to cell lysates in *rde-4+/+* animals, consistent with ALG-1 binding siRNAs and miRNAs with similar affinity (Figure 3E; Table S4)^32^.

Furthermore, unlike in GFP::RDE-1 co-IPs, there was no detectable difference in the enrichment of *nrfl-1* exogenous siRNAs and 23H-RNAs in HA::ALG-1 co-IPs from *rde-4-/-* compared to *rde-4+/+* animals (Figures 3F-3G; Table S4). Therefore, RDE-4 does not promote siRNA association with ALG-1. These results indicate that RDE-4 has a critical role in directing siRNA association with RDE-1 but not ALG-1. This role for RDE-4 likely ensures that siRNAs are preferentially paired with RDE-1 because of its ability to trigger high levels of 2° siRNAs from target mRNAs^38^.

Based on our earlier observation that siRNAs and 23H-RNAs are not depleted in *rde-1* mutants, these small RNAs do not require RDE-1 for stability. Therefore, the reduction in exogenous siRNA and 23H-RNA levels we observed in *rde-4* mutants is consistent with RDE-4 having a direct role in their biogenesis, rather than the decrease being an indirect consequence of RDE-1-dependent stability. This is supported by previous studies showing that RDE-4 is required for DCR-1 processing of siRNAs *in vitro*^10,13^. Together, these finding indicate that RDE-4 has dual functions in RNAi: first in facilitating DCR-1 processing of siRNAs, and second, in promoting their loading into RDE-1.

### Role of RDE-4 in 26G-RNA pathways

RDE-4 has also been implicated in the ALG-3/4 and ERGO-1 26G-RNA pathways, consistent with our earlier results showing widespread reductions in these small RNAs in *rde-4* mutants (Figures 1C-1F) ^6,7,42^. We therefore tested if ALG-3 and ERGO-1 depend on RDE-4 for 26G-RNA association. Despite a strong reduction in ALG-3/4 class 26G-RNAs in *rde-4-/-* compared to *rde-4+/+* cell lysates, their relative enrichment in GFP::ALG-3 co-IPs was unchanged (Figure 4A; Table S5). This indicates that while RDE-4 is essential for the formation or stability of ALG-3/4 class 26G-RNAs, it may not facilitate their binding with ALG-3. In contrast, both the levels and relative enrichment of ERGO-1 class 26G-RNAs in GFP::ERGO-1 co-IPs were strongly reduced in *rde-4-/-* compared to *rde-4+/+* animals, indicating that RDE-4 promotes 26G-RNA association with ERGO-1 (Figure 4B; Table S6).

**Figure 4.**
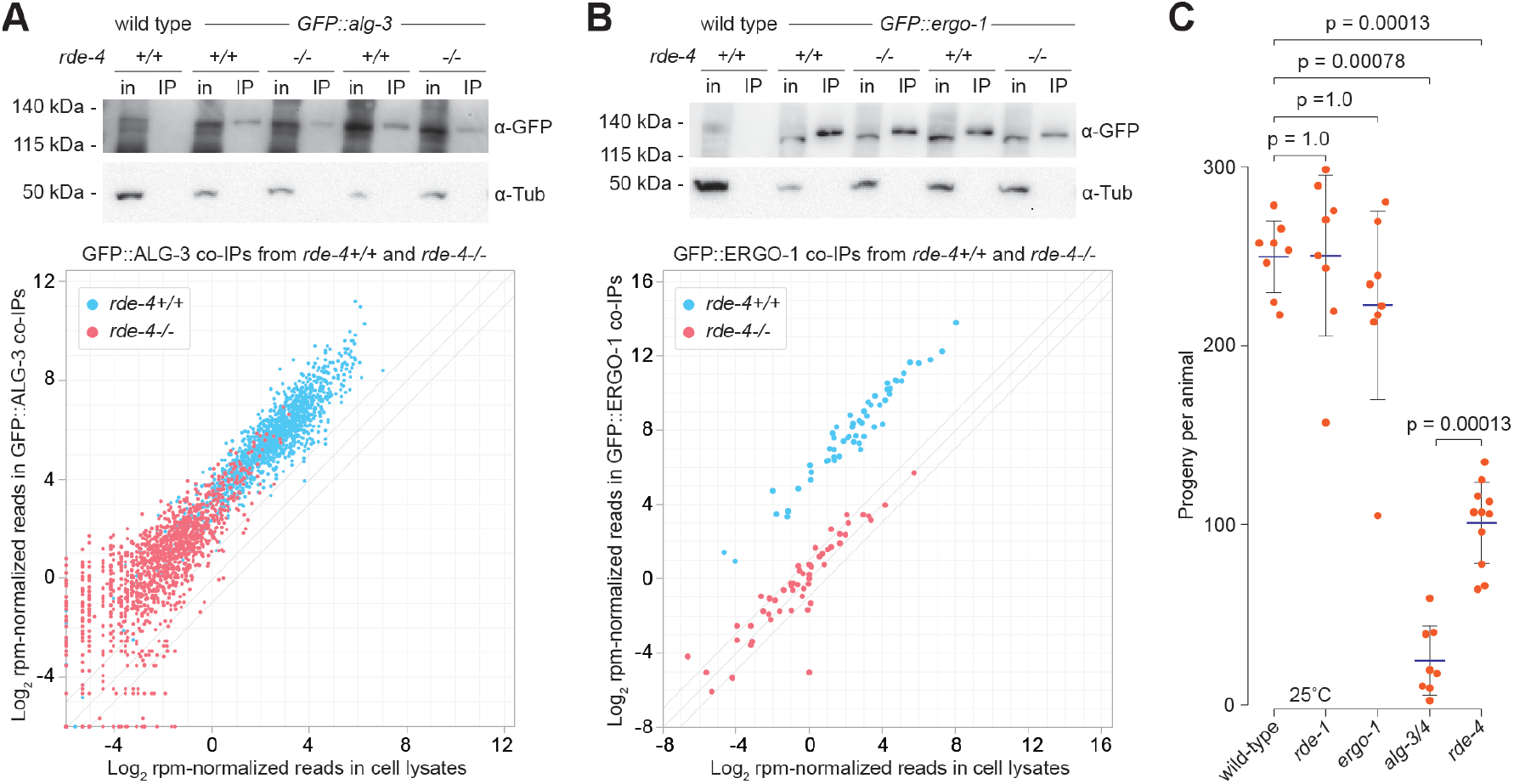
RDE-4 promotes 26G-RNA association with ERGO-1. **(A-B)** Scatter plots showing individual 26G-RNA features as the average log_2_ rpm-normalized sRNA-seq reads in cell lysates (x-axis) and in GFP::ALG-3 (**A**) or GFP::ERGO-1 co-IPs (**B**) (y-axis) from *rde-4+/+* or *rde-4-/-* animals. n=2 biological replicates. Western blot images above each scatter plot show GFP::ALG-3 (**A**) or GFP::ERGO-1 (**B**) levels in co-IP (IP) and cell lysate input (in) fractions. Tubulin is shown as a loading control. **(C)** Fertility of individual wild-type or *rde-1, ergo-1, alg-3/4*, or *rde-4* mutants grown at 25°C. Each point represents the total progeny from a single animal. Blue horizontal bars show means. Vertical bars show SD. *p*-values were calculated using the Mann-Whitney U test. n=8 (wild-type, *rde-1, ergo-1, alg-3/4*), or 11 (*rde-4*).

It is unclear why RDE-4 would be required for loading 26G-RNAs into ERGO-1 but not ALG-3. However, we and others have observed evidence that their immediate precursors are structurally distinct^43-46^. For example, ERGO-1 class 26G-RNA appear to derive from duplexes with presumed passenger strand sequences that are ∼19-20-nt long, which for reasons that are unclear match almost exclusively to the mRNA template and not the RdRP product^43^. Whereas the guide strand is almost exclusively antisense to the mRNA and thus produced from the RdRP transcript. The passenger strand is readily detectable in ERGO-1 co-IPs, comprising nearly 30% of 26G-RNA loci reads, suggesting that the 26G-RNA dsRNA duplex is loaded into ERGO-1 and the passenger strand is then discarded (Figures S3A-S3B). In ERGO-1 co-IPs from *rde-4-/-* animals, the passenger strand is still detectable, albeit at much lower levels, consistent with a role for RDE-4 in promoting the loading of the duplex into ERGO-1 (Figure S3C). In contrast, ALG-3 does not efficiently co-IP with sequences sense to the mRNA, with only ∼1% of 26G-RNA loci reads derived from the sense strand, corresponding to the hypothetical passenger strand (Figures S3D-S3F). Thus, we speculate that ALG-3 may load single-stranded 26G-RNAs, thereby bypassing the requirement for RDE-4 in siRNA loading, possibly utilizing a single-stranded RNA binding protein instead.

How might RDE-4 promote loading of small RNAs into Argonautes? One possibility is that it works similarly to R2D2 in *Drosophila*, which is proposed to position the siRNA duplex in the right orientation for loading into the Argonaute Ago2^30^. Another possibility is that RDE-4 functions similarly to the proposed, albeit questionable, role of the dsRNA-binding protein TRBP in mammals, acting as a bridge between Argonautes and Dicer to facilitate the formation of the RNA silencing complex^47,48^. This could indirectly influence loading by bringing the proteins into close proximity. RDE-4 has been shown to be in complexes with both DCR-1 and RDE-1, supporting either of these models^9,14,49^.

### RDE-4’s role in fertility likely relates to its function in the ALG-3/4 pathway

23H-RNAs and ERGO-1 class 26G-RNAs do not have clear roles in development, whereas ALG-3/4 class 26G-RNAs are required for proper sperm development^32,34,35,50-52^. RDE-4 is required for efficient exogenous RNAi, antiviral defense, proper chemotaxis, and optimal fertility at elevated temperatures^7,8,15,53,54^. Given our results demonstrating its global requirement in ALG-3/4 class 26G-RNA formation, we hypothesized that RDE-4’s role in fertility is related to its function within this pathway. Indeed, we observed a significant reduction in the mean number of progeny produced by *rde-4* mutants (Figure 4C). However, the total number of progeny produced by *rde-4* was still greater than that of *alg-3 alg-4* mutants likely because low levels of ALG-3/4 class 26G RNAs are still present and loaded into ALG-3, and likely also ALG-4, in *rde-4* mutants (Figure 4C). *ergo-1* and *rde-1* mutants did not have reduced brood sizes, suggesting that the lower numbers of progeny produced by *rde-4* mutants is due specifically to RDE-4’s role in the ALG-3/4 pathway (Figure 4C)^32,52^. Thus, RDE-4 has roles in antiviral defense, behavior, fertility, and development through its modular participation in multiple distinct small RNA pathways.

## Conclusions

*rde-4* was among the first RNAi factors identified by the Mello lab over twenty-five years ago^8,49^. These early studies pointed to a role for RDE-4 in siRNA biogenesis, which was later confirmed in elegant biochemical studies from the Bass lab^10,11,13^. However, our research uncovers a critical function for RDE-4 that was not evident in previous studies: its involvement in pairing siRNAs with the correct Argonautes. These findings highlight a dual role for RDE-4 in both exogenous and select endogenous siRNA pathways, broadening our understanding of its molecular functions and furthering our understanding of RNAi. In future studies, it will be important to identify the mechanism by which RDE-4 promotes Argonaute loading and whether other dsRNA-binding proteins fulfill this role in small RNA pathways where RDE-4 is not necessary.

## Supplemental Information

Document S1: Figures S1-S3. Related to Figures 2-4.

Tables S1-S4: Excel files containing geometric mean-normalized sRNA-seq counts and differential expression analysis. Related to Figures 1-3.

Tables S5-S6. Excel files containing rpm-normalized sRNA-seq counts. Related to Figure 4.

## RESOURCE AVAILABILITY

### Lead contact

Requests for additional information and resources should be directed to and will be fulfilled by the lead contact, Taiowa A. Montgomery.

### Material availability

The *rde-4* deletion allele generated in this study will be made available from the Caenorhabditis Genetics Center upon publication. Strains not distributed by the CGC are available on request.

### Data and code availability

Raw and processed high-throughput sequencing data generated in this study are available from the NCBI Gene Expression Omnibus (GEO; https://www.ncbi.nlm.nih.gov/geo/) under accession number GSE293782). The tinyRNA software used for analysis of sRNA-seq data is freely available from https://github.com/MontgomeryLab/tinyRNA.

## ACKNOWLEDGEMENTS

Thanks to Alivia Ball for help with media and solutions. Some strains used in this study were provided by the CGC, which is funded by the National Institutes of Health Office of Research Infrastructure Programs (P40 OD010440). This work was supported by the National Institutes of Health [R35GM119775 to T.A.M. and R35GM119656 to C.M.P.].

## METHODS

### Strains

WM45[*rde-1*(ne300) V], TAM151[*rde-1(ram40)* V] ,WM158[*ergo-1(tm1860)* V], WM300[*alg-4(ok1041)* III; *alg-3(tm1155)* IV], JMC205[*alg-3(tor141[GFP::3xFLAG::alg-3])* IV], JMC211[*ergo-1(tor147[GFP::3xFLAG::ergo-1a])* V], TAM10[*unc-119(ed3); ram4([pCMP2]alg-1::3XHA::TEV::alg-1 + Cbr-unc-119(+))*], and USC1080[*rde-1(cmp133[(gfp + loxP + 3xFLAG)::rde-1])* V] were previously described^8,32,35,41,51,55,56^. PHX6665[*rde-4(syb6665)* III] was generated by SunyBiotech (Fuzhou, China) using CRISPR/Cas9 genome editing^57-60^. TAM122[*rde-4(syb6665)* III*; ram4([pCMP2]alg-1::3XHA::TEV::alg-1 + Cbr-unc-119(+))* IV] was generated by crossing PHX6665 with TAM10. TAM98[*rde-4(syb6665)* III*; rde-1(cmp133[(gfp + loxP + 3xFLAG)::rde-1])* V] was generated by crossing USC1080 to PHX6665. TAM104[*rde-4(syb6665)* III*; alg-3(tor141[GFP::3xFLAG::alg-3])* IV] was generated by crossing JMC205 to PHX6665. TAM106[*rde-4(syb6665)* III*; ergo-1(tor147[GFP::3xFLAG::ergo-1a])* V] was generated by crossing JMC211 to PHX6665. USC1557[*dcr-1(cmp321[dcr-1::2xHA])* III] and USC1556[*rde-4(cmp320[rde-4::2xFLAG])* III] were generated via CRISPR-Cas9 genome editing in wild-type animals by injecting purified Cas9 protein along with guide RNAs and repair templates^61^. For USC1556, the guide RNA sequence was: /AltR1/rUrUrUrCrArArCrArCrCrUrArUrGrArUrUrUrCrArGrUrUrUrUrArGrArGrCrUrArUrGrCrU/AltR2/. The repair template sequence was: tataatcattcagacgcttcatttttcaggaatacgcaataatatttTCATTTGTCATCATCGTCTTTATAATCCTTATCGTCGTCATCCTTGTAGTCATCgGTGAAATCATAGGTGTTGAAATGGATAATCGCCGATTTACAAGCACACT GTTTA. For CMP1557, the guide RNA sequence was: /AltR1/rUrUrUrUrCrArGrArCrCrArArUrArArUrGrGrUrCrGrUrUrUrUrArGrArGrCrUrArUrGrCrU/AltR2/. The repair template sequence was: attgtaatttttgaacattatcaattttccctgttttcagaccaataATGTATCCTTATGATGTACCTGATTATGCCTACCCATACGACGTTCCAGACTACGCTGTCAGGGTAAGAGCTGATTTACAATGTTTTAACCCCAGGGACTACC AGgt.

### Animal growth conditions

Animals were cultured at 20°C on NGM plates (3 g/l NaCl, 17 g/l agar, 2.5 g/l peptone, 1 mM CaCl_2_, 5 µg/ml cholesterol, 1 mM MgSO_4_, and 25 mM KPO_4_ buffer) seeded with 2 mL (10 cm plates) or 0.5 mL (6 cm plates) of *E. coli* OP50 culture. The bacteria was grown overnight at 37°C in LB Broth (NaCl, 5 g/L, Tryptone, 10 g/L, Yeast Extract, 5 g/L – Sigma, cat# L3022), then applied to plates and allowed to air dry with lids closed for approximately 3-4 days to allow the formation of a bacterial lawn. The NGM was supplemented with 1% Nystatin and 2.5% Streptomycin to prevent contamination.

### Stage synchronization

To obtain embryos, gravid adult hermaphrodites were collected from NGM plates by washing with M9 buffer into a 15 mL conical tube. After removing excess buffer, leaving 1.5 mL, a bleaching solution (2.5 mL of 1 M NaOH and 1 mL of 5% sodium hypochlorite) was added to lyse hermaphrodites while preserving the eggs. Samples were vortexed for ∼10 minutes, with periodic checks to ensure disintegration of carcasses. Once the solution consisted of mostly embryos, samples were centrifuged at 1.9 krcf for 30 seconds, and the supernatant was carefully discarded without disturbing the egg pellet. The pellet was washed three times by resuspension in 15 mL of M9 followed by centrifugation at 1.9 krcf for 30 seconds. Synchronized L1-arrested larvae were obtained by incubating the isolated embryos in 2 mL of M9 buffer (composition detailed below) at 15°C with gentle rotation (∼15 rpm) for 72 hours.

### Protein structure analysis

Structures of *Drosophila* R2D2 and Loqs-PD and of *C. elegans* RDE-4 were predicted using AlphaFold3 with protein sequences obtained from UniProt^62,63^. The structures were analyzed and images generated using ChimeraX^64^.

### RNA isolation

To extract RNA from whole animals at either gravid adult stage (72 hours at 20°C following L1 synchronization) or L4 larval stage (52 hours at 20°C post-synchronization) animals were harvested from plates, washed three times with M9 buffer, and immediately flash-frozen in liquid nitrogen. Co-IPs and corresponding cell lysates were used directly for RNA extraction. RNA isolation was performed using TRIzol reagent (Life Technologies, cat# 15596018) following the manufacturer’s protocol, with the addition of a second chloroform extraction step to improve purity.

### sRNA-seq

RNA ranging from ∼16 to 30 nucleotides was size-selected by gel extraction using 17% polyacrylamide/urea gels, from RNA, in some instances pre-treated with RNA 5′ pyrophosphohydrolase (New England Biolabs, cat# M0356S) to convert 5′ triphosphates to 5’ monophosphates (as noted in figure legends), which enhances ligation efficiency of 22G-RNAs^65^. Library preparation was performed using the NEBNext Multiplex Small RNA Library Prep Set for Illumina (NEB, cat# E7300S), following the manufacturer’s instructions, except that the 3′ ligation step was carried out at 16°C for 18 hours to optimize recovery of methylated small RNAs. PCR amplified small RNA libraries were size-selected on 10% polyacrylamide non-denaturing gels, and sequencing was conducted on an Illumina HiSeq X or NovaSeq X Plus (PE150) by Novogene. Only forward strand reads were retained for analysis.

### sRNA-seq data analysis

Small RNA sequencing data was processed and visualized using the tinyRNA pipeline with default settings^66-69^. Analyses were conducted using the *C. elegans* WS279 reference genome sequences and annotations^33^. Annotations for 23H-RNAs and other small RNA classes were previously published in GFF3 format^32^. Computing of normalized counts using the geometric mean method and statistical analysis using the Wald test were performed within the tinyRNA framework using the DESeq2 R package^70^. Additional plotting and statistical analyses were carried out using Matplotlib, R, IGV, and Adobe Illustrator^71-73^.

### RNAi

Synchronized L1 larvae were plated on RNAi plates (NGM supplemented with IPTG [1.2 mg/mL] and Carbenicillin [25 ug/mL]) seeded with *E. coli* HT115 bacteria expressing dsRNA matching *dcr-1, nrfl-1, oma-1*, or empty vector (L4440)^74^. Animals were grown for 72 hours at 20°C and gravid adults were collected for protein and RNA isolation, where applicable. For HA::DCR-1 protein isolation, animals were flash frozen in liquid nitrogen and then ground using mortars and pestles, following the same protocol as for Co-IPs.

### Co-IPs

Co-IPs were performed for GFP::3xFLAG::RDE-1, 3xHA::TEV::ALG-1, GFP::3xFLAG::ALG-3, and GFP::3xFLAG::ERGO-1 from 2-3 biological replicates, each containing ∼12,000 gravid adult *C. elegans* grown for 52 hours (L4 larvae for GFP::3xFLAG::ALG-3 co-IPs) or 72 hours (adults, all other co-IPs) after L1 synchronization. Animals were harvested from NGM plates to 15 mL tubes using M9 buffer (3 g/l KH_2_PO_4_, 6 g/l Na_2_HPO_4_, 5 g/l NaCl, 1 mM MgSO_4_) and washed three times with ∼10 mL of M9 buffer to remove bacteria. The buffer was removed, and 1.2 mL of lysis buffer was added (50 mM Tris-Cl pH 8.0, 100 mM KCl, 2.5 mM MgCl_2_, 0.1% Igepal CA-630, and 1X protease inhibitor cocktail - Pierce, cat# 88266). The solution was flash-frozen when transferred to a mortar with liquid nitrogen and ground using pestles. Lysates were transferred to 1.5 mL tubes and clarified by centrifugation at 12,000 × g for 10 minutes at 4°C. Supernatants were divided into input and co-IP fractions. For co-IP, 25 µL of GFP-Trap Magnetic Agarose Beads (ChromoTek, Proteintech, cat# gtma-100) or HA Affinity Matrix (Roche, cat# 11815016001) was incubated with the lysates (1 mL) for 1 hour at 4°C while rotating. Beads were then separated on a magnetic rack or by centrifugation at 500 rcf and washed three times with 1 mL of lysis buffer. Following 3 washes, beads were split between protein and RNA fractions. For protein fractions, samples were denatured by heating at 95°C for 5 minutes in lysis buffer with 1×Blue Protein Loading Dye (New England Biolabs, cat# B7703S: 62.5 mM Tris-HCl (pH 6.8), 2% (w/v) SDS, 10% glycerol, 0.01% (w/v) bromophenol blue) supplemented with 50 mM DTT. RNA fractions were treated as described above.

### qRT-PCR

qRT-PCR was performed using a custom TaqMan Gene Expression Assay for an F43E2.6 23H-RNA (Life Technologies, cat# 4331348, target sequence: UUUGCCGAUGUUUCUGAGAUGUC) or miR-1 (Life Technologies, assay name: hsa-miR-1, cat# 4427975) according to the manufacturer’s protocol. Expression levels of the F43E2.6 23H-RNA and miR-1 were measured on a Bio-Rad CFX96 Real-Time PCR Detection System. Ct values were averaged from three technical replicates for each of three independent biological samples. Relative quantification of the F43E2.6 23H-RNA normalized to miR-1 was determined using the 2^-ΔΔCt method^75^. Plotting and statistics were conducted in Microsoft Excel and GraphPad Prism.

### Western blots

Protein samples from co-IPs and their corresponding cell lysates were separated on 4-12% Bolt Bis-Tris Plus 15-well gels (Invitrogen, cat# NW04125BOX). After separation, proteins were transferred to nitrocellulose membranes and probed with anti-GFP (Santa Cruz Biotechnology, cat# sc-9996 HRP; 1:100), anti-HA (Roche, distributed by Merck, cat# 12013819001 HRP; 1:500), and anti-tubulin (Abcam, cat# ab40742 HRP; 1:1000) antibodies. Blots were imaged using a FluorChem E Imaging System (ProteinSimple).

### Fertility

Fertility was assessed in animals grown since conception at 25°C. These animals were obtained by transferring their parents from the maintenance temperature of 20°C to 25°C at the L4 stage, before the onset of reproduction. The animals grown at 25°C were singled out onto individual plates at the L4 stage and transferred daily to new plates to allow for a clear distinction between the adult parent and the developing larval stage progeny. After the parent was removed from a plate, the number of L2-L3 larval stage progeny was counted on a stereoscope. The total number of progeny produced by each animal across its reproductive span was summed. The experiment was ended when no viable progeny were produced for 24 hours.

### Statistics

Mann-Whitney U tests were used to calculate *p*-values using GraphPad Prism when evaluating the fertility assay data. Two-sample t-tests were applied to qRT-PCR, Western blot quantifications, and comparisons between co-IP and corresponding input fractions, with calculations performed in Microsoft Excel. When multiple comparisons were made, *p*-values were adjusted using the Bonferroni correction.

