## Supplemental Figures for "Argonaute-siRNA loading via the RNA-binding protein RDE-4 in *C. elegans*"

**Figures S1-S3.**

### Supplemental Figures

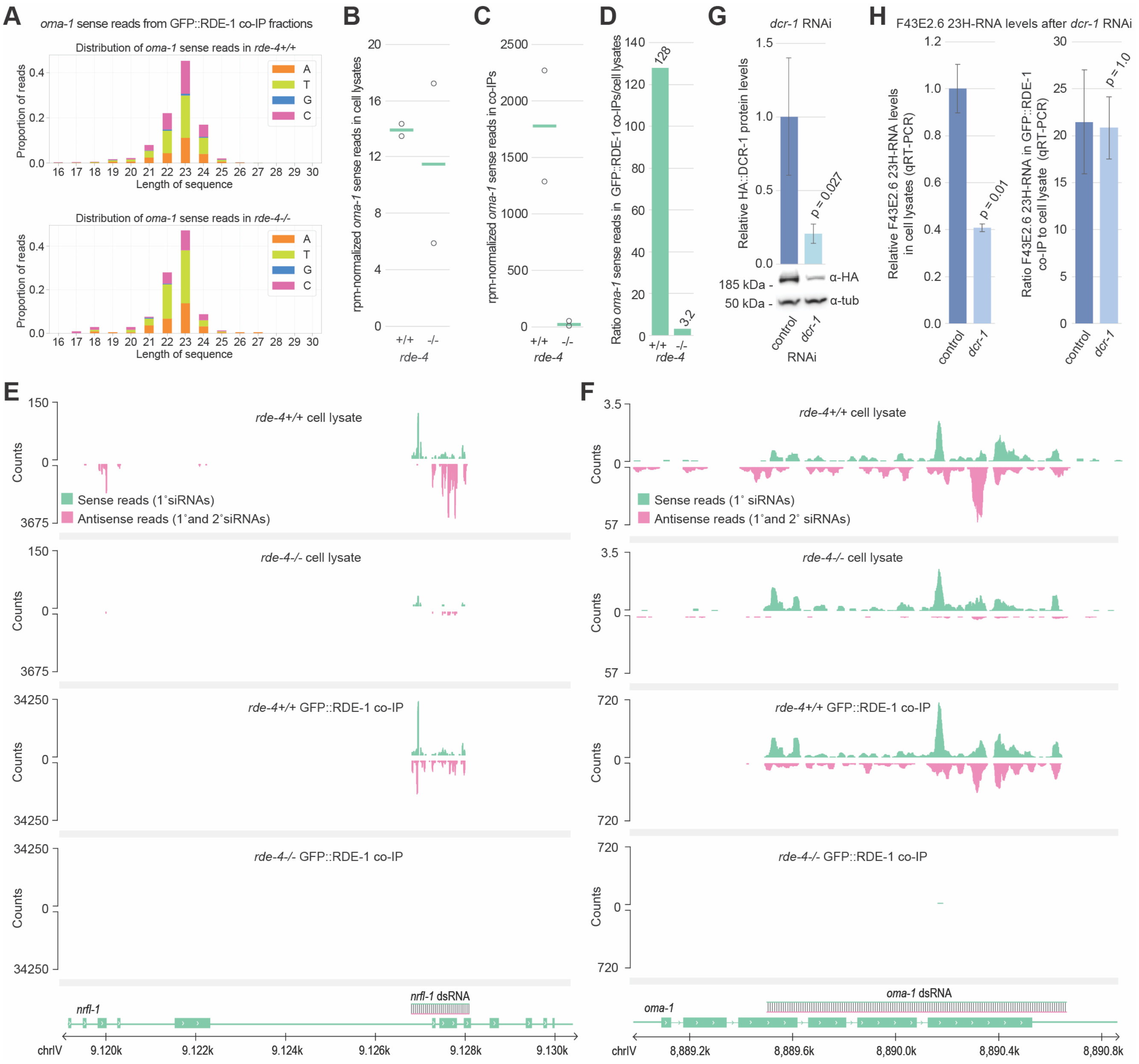

**Figure S1. RDE-4 promotes the association of exogenous primary siRNAs with RDE-1. (A)** Size distribution and 5′-nt identity of *oma-1* sense siRNAs from GFP::RDE-1 co-IPs from *rde-4+/+* and *rde-4-/-* gravid adults following *nrfl-1* and *oma-1* RNAi (only *oma-1* is shown). Data from one of two biological replicates are shown. (**B-C**) Total rpm-normalized counts for *oma-1* sense siRNAs in cell lysates (**B**) and GFP::RDE-1 co-IPs (**C**). Green lines represent the mean from two biological replicates; individual replicate values are shown as circles. **(D)** Enrichment of total rpm-normalized *oma-1* sense siRNAs reads in GFP::RDE-1 co-IP libraries relative to corresponding cell lysates (data as in A-C). **(E-F)** siRNA read distribution across the sense and anti-sense strands of *nrfl-1* (E) and *oma-1* (F) from cell lysates and GFP::RDE-1 co-IPs from *rde-4+/+* and *rde-4-/-* gravid adults. Data from one of two biological replicates is shown. **(G)** Western blot analysis of HA::DCR-1 levels in gravid adults treated with control (empty L4440 vector) or *dcr-1* RNAi. Tubulin serves as a loading control. Data from one of three biological replicates is shown. The graph shows average HA::DCR-1 levels quantified from the blots. The error bars represent SD of the mean from three biological replicates. The *p*-value was calculated using a two-sample t-test. **(H)** Average relative levels of the most abundant F43E2.6 23H-RNA normalized to miR-1 levels, as measured by qRT-PCR in cell lysates (left) and enrichment in GFP::RDE-1 co-IPs relative to cell lysates (right) from wild-type gravid adults treated with control (L4440) or *dcr-1* RNAi. Error bars represent the SD from the mean of three biological replicates. *p*-values were calculated using two-sample t-tests.

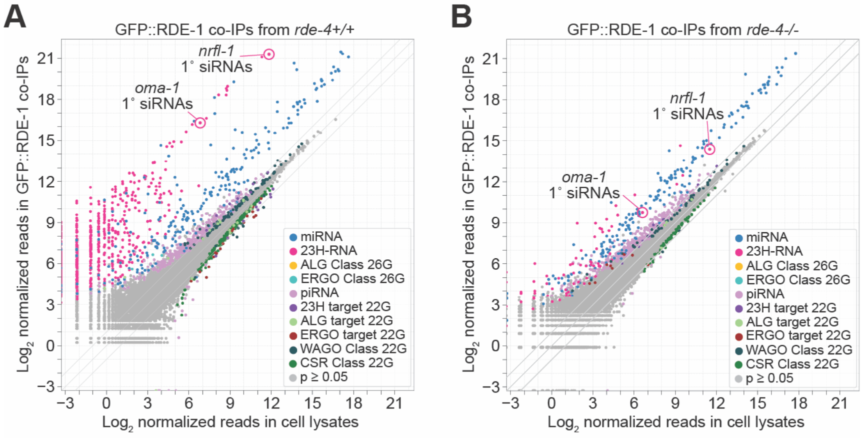


**Figure S2. RDE-1-small RNA interactions in *rde-4* mutants. (A-B)** Scatter plots showing individual small RNA features as the average $\log_2$ geometric mean-normalized sRNA-seq reads from two biological replicates in cell lysates (x-axis) and GFP::RDE-1 co-IPs (y-axis) from *rde-4*+/+ (A) and *rde-4-/-* (B) gravid adults. Small RNA classes are color-coded. Exogenous siRNAs mapping to *nrfl-1* and *oma-1* are circled.

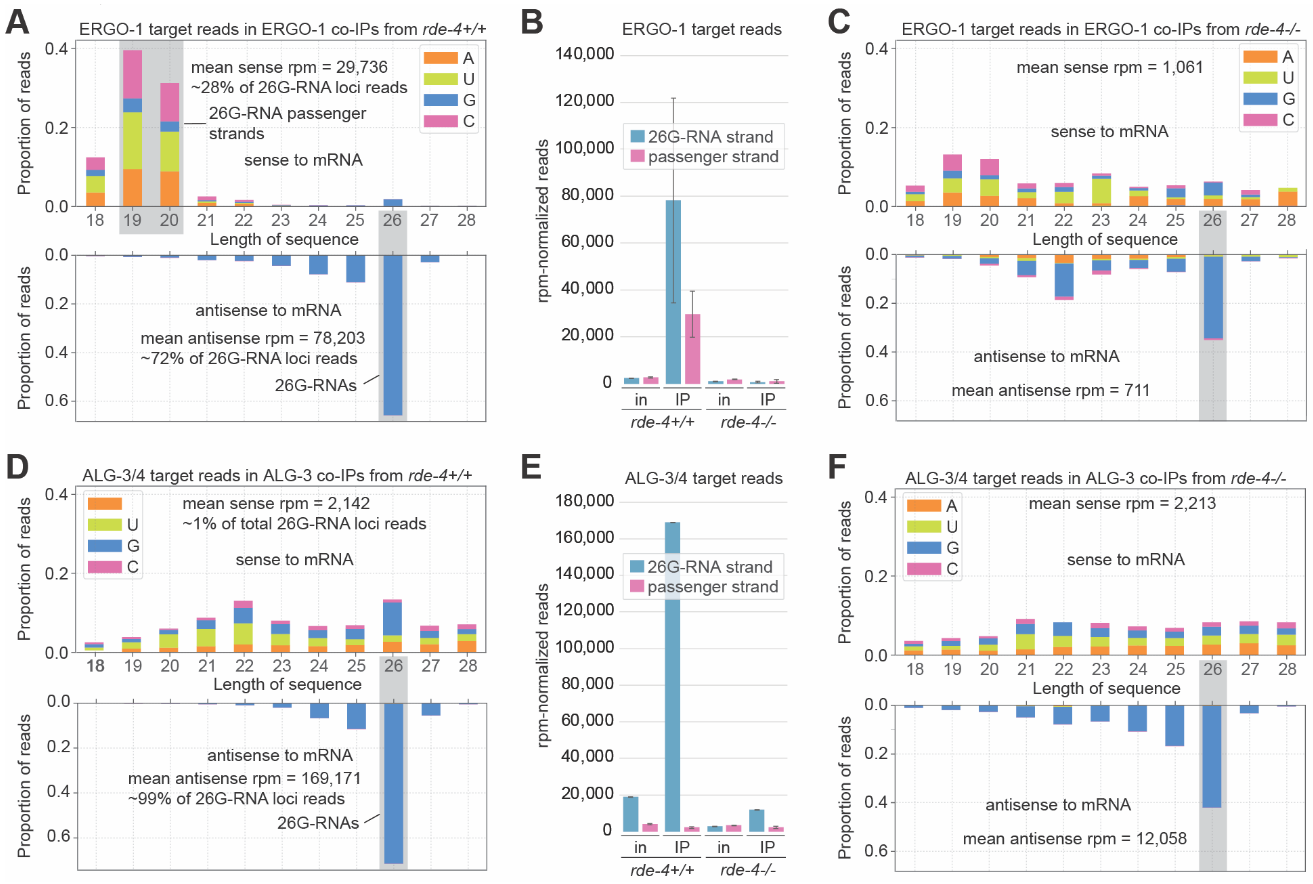


**Figure S3. 26G-RNA duplex strand analysis from GFP::ERGO-1 and GFP::ALG-3 co-IPs. (A)** Size distribution and 5′-nt identity of small RNA reads mapping to genes targeted by ERGO-1 class 26G-RNAs from ERGO-1 co-IPs from *rde-4+/+* gravid adults. The top graph represents sense reads relative to the corresponding mRNA (passenger strand reads) and the bottom graph shows antisense reads (guide strand reads). Data from one of two biological replicates is shown. **(B)** Average rpm-normalized ERGO-1 target sRNA-seq reads from cell lysates and ERGO-1 co-IPs from *rde-4+/+ and rde-4-/-* gravid adults (data as in A). Error bars represent SD from the mean of two biological replicates. **(C)** Size distribution and 5′-nt identity of small RNA reads mapping to genes targeted by ERGO-1 class 26G-RNAs from ERGO-1 co-IPs from *rde-4-/-* gravid adults (same data as in B). **(D)** Same as in (A) but for sRNA-seq reads from cell lysates and ALG-3 co-IPs from *rde-4+/+* L4 larvae. Data from one of two biological replicates is shown. **(E)** Average rpm-normalized ALG-3/4 target sRNA-seq reads from cell lysates and ALG-3 co-IPs from *rde-4+/+* and *rde-4-/-* gravid adults (data as in D). Error bars represent SD from the mean of two biological replicates. (**F**) Same as in (C) but for sRNA-seq reads from cell lysates and ALG-3 co-IPs from *rde-4-/-* L4 larvae (same data as in E). Data from one of two biological replicates is shown.
